# *De novo* assembly of the *Pasteuria penetrans* genome reveals high plasticity, host dependency, and BclA-like collagens

**DOI:** 10.1101/485748

**Authors:** Jamie N Orr, Tim H Mauchline, Peter J Cock, Vivian C Blok, Keith G Davies

## Abstract

*Pasteuria penetrans* is a gram-positive endospore forming bacterial parasite of *Meloidogyne* spp. the most economically damaging genus of plant parasitic nematodes globally. The obligate antagonistic nature of *P. penetrans* makes it an attractive candidate biological control agent. However, deployment of *P. penetrans* for this purpose is inhibited by a lack of understanding of its metabolism and the molecular mechanics underpinning parasitism of the host, in particular the initial attachment of the endospore to the nematode cuticle. Several attempts to assemble the genomes of species within this genus have been unsuccessful. Primarily this is due to the obligate parasitic nature of the bacterium which makes obtaining genomic DNA of sufficient quantity and quality which is free from contamination challenging. Taking advantage of recent developments in whole genome amplification, long read sequencing platforms, and assembly algorithms, we have developed a protocol to generate large quantities of high molecular weight genomic DNA from a small number of purified endospores. We demonstrate this method via genomic assembly of *P. penetrans*. This assembly reveals a reduced genome of 2.64Mbp estimated to represent 86% of the complete sequence; its reduced metabolism reflects widespread reliance on the host and possibly associated organisms. Additionally, apparent expansion of transposases and prediction of partial competence pathways suggest a high degree of genomic plasticity. Phylogenetic analysis places our sequence within the Bacilli, and most closely related to *Thermoactinomyces* species. Seventeen predicted BclA-like proteins are identified which may be involved in the determination of attachment specificity. This resource may be used to develop *in vitro* culture methods and to investigate the genetic and molecular basis of attachment specificity.

## 2. DATA SUMMARY

1. *Pasteuria penetrans* RES148 genome has been deposited in the European Nucleotide Archive; accession number: ERZ690503
2. PacBio reads in ENA, accession ERR2736894 and ERR2736893
3. Legacy Illumina reads in ENA, accession ERR2736890
4. Scripts used in this analysis can be accessed on GitHub: https://github.com/BioJNO/Ppenetrans_genomics

I/We confirm all supporting data, code and protocols have been provided within the article or through supplementary data files.

## 3. IMPACT STATEMENT

*Pasteuria penetrans* is a natural bacterial antagonist to the most economically damaging nematodes in agriculture. It may be possible to reduce or replace the use of rapidly declining chemical nematicides with biological control using this organism. However, this bacterium has high host specificity and is extremely difficult to mass produce. To provide a resource which is likely to help to solve these issues, we have generated the first genomic assembly of any bacterium within this genus. The genomic assembly generated is small but near complete, reflecting reliance on the host for many metabolic processes. This provides key insights into the metabolism of this bacterium which are likely to be of significant commercial and scientific interest. We have identified proteins which may be directly involved in host specificity, of interest to researchers involved in both agricultural and evolutionary biology. The methods we describe may be used to vastly expand the current availability of genomic data within this genus of bacteria, and may be applicable to other challenging genomic sequencing projects.

## 4. INTRODUCTION

*Pasteuria penetrans* is an endospore forming Firmicute which is an obligate parasite of root-knot nematode (RKN, *Meloidogyne* spp.), a globally distributed genus of plant parasitic nematodes which are among the most economically devastating in agriculture [1, 2]. *P. penetrans* act as natural antagonists to RKN via two key mechanisms. Firstly, attachment of endospores to the nematode cuticle hinders movement, migration through the soil, and thus root invasion [3, 4]. Secondly, bacterial infection of the plant feeding nematode results in sterilisation. As such *P. penetrans* is of considerable interest as a biological alternative to chemical nematicides. The effective application of *Pasteuria* spp. for this purpose is currently limited by lack of understanding of nematode attachment specificity and *in vitro* culture method development. Attachment of the endospore to the nematode cuticle is a determinative process in infection [5]. *Pasteuria* spp. may exhibit extremely fastidious attachment profiles including species and population specificity [6, 7]. Attempts to characterise the molecular basis of attachment have identified two components which appear to be involved in this process from the perspective of the endospore: collagen and N-acetyl-glucosamine (NAG). Treatment of endospores with collagenase, NAGase, and the collagen binding domain of fibronectin inhibit attachment [8–11]. This has prompted the current “Velcro-model” of attachment involving bacterial collagen-like fibres, observable under electron microscopy on the exosporium surface, and nematode cuticle associated mucins [12]. Recently, Phani *et al.* [13] demonstrated that knockdown of a mucin like gene, *Mi-muc-1*, reduced cuticular attachment of *P. penetrans* endospores to *M. incognita*. However, the exact nature of this host-parasite interaction is not known at the genetic or molecular level. Additionally, no published medium is available *in vitro* production of *P. penetrans* [14]. This is attributable to its obligate parasitism of nematodes that are themselves obligate parasites. In short, it is not yet known what *P. penetrans* requires from its host in order to proliferate. Adding to this complex picture is the apparent influence of “helper-bacteria” which have been implicated in growth promotion [15]. This means that in order to complete its life cycle *P. penetrans* may rely on metabolic and/or signalling pathways from the plant, the nematode, and from associated bacteria.

The difficulty of obtaining genomic DNA of sufficient quantity, quality, and purity from *P. penetrans* has so far impeded attempts to obtain a high quality genomic assembly. An assembly of 20,360bp with an N50 of 949bp (GCA_000190395.1) from *P. nishizawae*, a bacterial pathogen infective of the soybean cyst nematode (*Heterodera glycines)* [16], is available, along with some PCR generated marker gene sequences, and a 2.4Mbp Sanger shotgun sequence generated genome survey sequence of *P. penetrans* [17]. A 2.5Mbp shotgun sequence assembly in 563 contigs with a GC content of 48.3% was described but not published [18]. Recent advances in both whole genome amplification (WGA) technology and assembly algorithms have enabled genomic assembly from low abundance microorganisms, single cells, and complex samples [19–24]. In order to provide insights into the metabolism and attachment specificity of *P. penetrans* we attempted to generate genomic assemblies of strain RES148 using two data sets. First we attempted to improve assembly metrics of previously generated Illumina data using GC-coverage plots to visualise, identify, and remove contamination [25]. Second, we developed a simple method of purifying small numbers of *P. penetrans* endospores. Then, using multiple displacement amplification (MDA), we were able to generate genomic DNA of sufficient length and quantity for PacBio sequencing and *de novo* assembly of this strain. Assembled genomic data reveals a reduced genome with a reduced metabolism, and unusually high plasticity. Several predicted proteins which may be involved in the attachment of endospores to the nematode cuticle are also identified. When compared to published and described genomic data this sequence presents a significant improvement in all available metrics.

## 5. METHODS

### 5.1 Strains and culture

*Pasteuria penetrans* were cultivated on *Meloidogyne javanica* 16, isolated from Greece and kindly provided by Emmanuel Tzortzakakis. *Pasteuria penetrans* strain RES148 is a highly passaged strain derived from strain RES147 (also referred to as strain PNG in earlier literature) which was isolated from Papua New Guinean soils by Dr John Bridge. Approximately 5000 juvenile *M. javanica* were encumbered with 3-5 endospores by centrifugation at 9500g for two minutes as described by Hewlett and Dickson [26]. Juvenile nematodes were counted and assessed visually for the attachment of spores under an inverted microscope (Hund Wilovert^®^) at 200x magnification. Spore encumbered juveniles were re-suspended in 5ml of sterile distilled water and inoculated onto 4-week old tomato plants (cv. MoneyMaker) in peat and allowed to grow at 25°C for 900 growing degree days. Roots were washed thoroughly in tap water to remove soil and stored at −20°C.s

### 5.2 Removal of contamination

Tomato roots containing *P. penetrans* infected *M. javanica* were subjected to three freeze-thaw cycles to weaken root tissue. Approximately 100 *P. penetrans* infected *M. javanica* females were dissected from root material in sterile 1X PBS solution using mounted needle and forceps. Dissected females were transferred to a clean 1.5ml LoBind Eppendorf (Sigma) and washed three times in 1ml of HPLC water containing Triton X-100 0.5%. Washed females were burst with a micropestle in a clean 1.5ml LoBind Eppendorf (Sigma), and the contents subjected to a series of washes at room temperature: first three times in 1ml HPLC water; second three times in 1ml 70% ethanol; and finally, once in a 500μl 0.05% sodium hypochlorite solution, before density selection on a sterile 1.25g/ml sucrose gradient. All centrifugation steps were at 20817g for 15 minutes except spore pelleting after sodium hypochlorite incubation which was 5 minutes. The resulting clean endospore suspensions were inspected at 1000x magnification (Zeiss Axiosop).

### 5.3 DNA extraction and MDA

Clean endospores were subjected to a 30-minute lysozyme digestion, spun to pellet, ground for 1 minute with a micropestle, re-suspended in 4μl scPBS, and then passed immediately into the Repli-g whole genome amplification protocol for single cells (Qiagen). A 16hr isothermal amplification protocol produced 15μg of genomic DNA. Amplified genomic material was visualised on a 0.5% agarose gel, quantified with a Qubit hsDNA quantification kit (ThermoFisher), and assayed for *Pasteuria* spp. specific 16S rRNA gene sequence using primers 39F and 1166R as previously described [27]. The resultant library was submitted to Oslo Genomic Sequencing Centre for two runs of PacBio SMRT cell sequencing. Legacy WGA Illumina data, from the same strain was included in these analyses. The clean-up protocol for this material has been previously described [27]. The WGA, debranching, S1 nuclease treatment, and Illumina library prep for this sample are described in supplementary methods file 1.

### 5.4 Legacy Illumina assembly

Legacy illumina data was reduced to exclude contaminating material using the BlobTools pipeline [25]. Briefly, Illumina data was assembled using MIRA (v4.9.6) [28]; contigs were aligned against the NCBI non-redundant protein database (nr, circa June 2015) using BLAST (v2.7.1) [29]; raw illumina reads were mapped to contigs using BWA (v0.7.12) [30]; GC coverage plots were generated using BlobTools (v1.0) [25]; reads mapping to contaminant contigs were removed using mirabait (v4.9.6); and “clean” Illumina read sets were re-assembled using MIRA. This was repeated iteratively, a total of 14 times, until no further improvements in assembly metrics were observed.

### 5.5 PacBio assembly

The PacBio sequence reads were trimmed and assembled initially using Canu (v1.5) [3], this initial assembly was polished to correct sequencing errors twice, first using FinisherSC (v2.1) [31], and then using Arrow (v2.1.0) with raw read alignment from PBalign (v0.3.0), both from PacBio’s SMRT^®^ Analysis suite (v4.0.0). Hybrid Pacbio and Illumina assemblies were compiled using Spades (v3.5.0) [20]. Assembly merging was carried out using quickmerge (v0.2) [32].

### 5.6 Genome quality assessment

Genome completeness, heterogeneity, and contamination were determined by alignment with lineage specific marker genes using CheckM (v1.0.7) [11]. Genomic average nucleotide alignments were carried out using Pyani (v0.2.7) using mummer (ANIm) [33].

### 5.7 Comparative genomics

Coding sequences were predicted from *P. penetrans* PacBio assemblies using RASTtk (http://rast.nmpdr.org, 18-07-2017). A multiple gene maximum likelihood tree was generated using bcgTree (v1.0.10) [34] with alignment to related genomic annotations from assemblies listed in the data bibliography. Metabolic modelling of *P. penetrans* was carried out using BLASTKoala [35] to identify KEGG database orthologues and KEGG Mapper (v3.1) to construct and visualise pathways on the KEGG server [36]. Metabolic profiles of *Thermoactinomyces vulgaris* (GCA_001294365.1), *Xiphinematobacter* spp. (GCA_001318295.1) and the *Wolbachia* symbiont of *Brugia malayi* (GCA_000008385.1) were also generated for comparison. Coding sequences were re-predicted from *B. subtilis*, *B. thuringiensis*, *B. cereus*, *T. vulgaris*, and *C. difficile* (Genbank accessions 6,9,16,17, and 20 as listed in the data bibliography) using RASTtk. Resultant predicted proteomes were clustered with *P. penetrans* predicted proteins using OrthoFinder (v2.2.1) [37], and functionally annotated using InterProScan (v5.29-68.0) [38]. Clusters and annotations were aggregated using KinFin (v1.0) [39]. Clusters annotated with sporulation related terms were extracted using an R script (this study). Cluster intersections were plotted using UpSetR (v1.3.3) [40].

### 5.8 Putative attachment proteins

*Pasteuria penetrans* gene predictions were interrogated for collagens using Pfam collagen domain (PF01391) and HMMER (v3.1b2) hmmsearch (http://hmmer.org/). Predicted collagens were aligned to contigs and to each other using BLASTn (v2.7.1) [13]. Unique collagen sequence structural models and predicted binding sites were produced using the RaptorX server [41]. Surface electrostatic potential was computed using the Adaptive Poisson-Boltzmann Solver (APBS) [42, 43]. RaptorX generated protein pdb files were converted to pqr using pdb2pqr [44]. Command line apbs (v1.5) was used to generate electrostatic potential maps. Protein structure visualisations were generated in the NGL viewer [45, 46]. BclA-like signal peptide prediction was performed with PRED-TAT [47].

## 6. RESULTS

### 6.1 Assembly

The highest scoring PacBio assembly resulted from a 200x coverage assembly of the second SMRTcell only. This assembly consisted of five contigs, with a total size of approximately 2.64Mbp, an N50 of 2.26Mbp and a completeness score of 86%. Some coding sequences were predicted in one version of the assembly but not in another, including lineage specific marker genes used by CheckM for completeness scoring. A high number of contaminant and heterogenic markers were observed in the legacy Illumina data assemblies; however, this was significantly reduced using the BlobTools pipeline (Fig. 1 and Fig 2). BLAST annotation of lineage specific marker genes returned by CheckM within raw and cleaned Illumina assemblies returned with 73% and 74% of markers aligning to *Pelosinus* spp. with an average identity of 93% and 92% respectively. *Clostridium* spp. returned as the best hit in 14% of markers in both Illumina assemblies with an average identity of 89%. Although the GSS also scored highly for contamination no high scoring BLAST hits indicating specific identifiable contaminants were observable.

**Figure 1:**
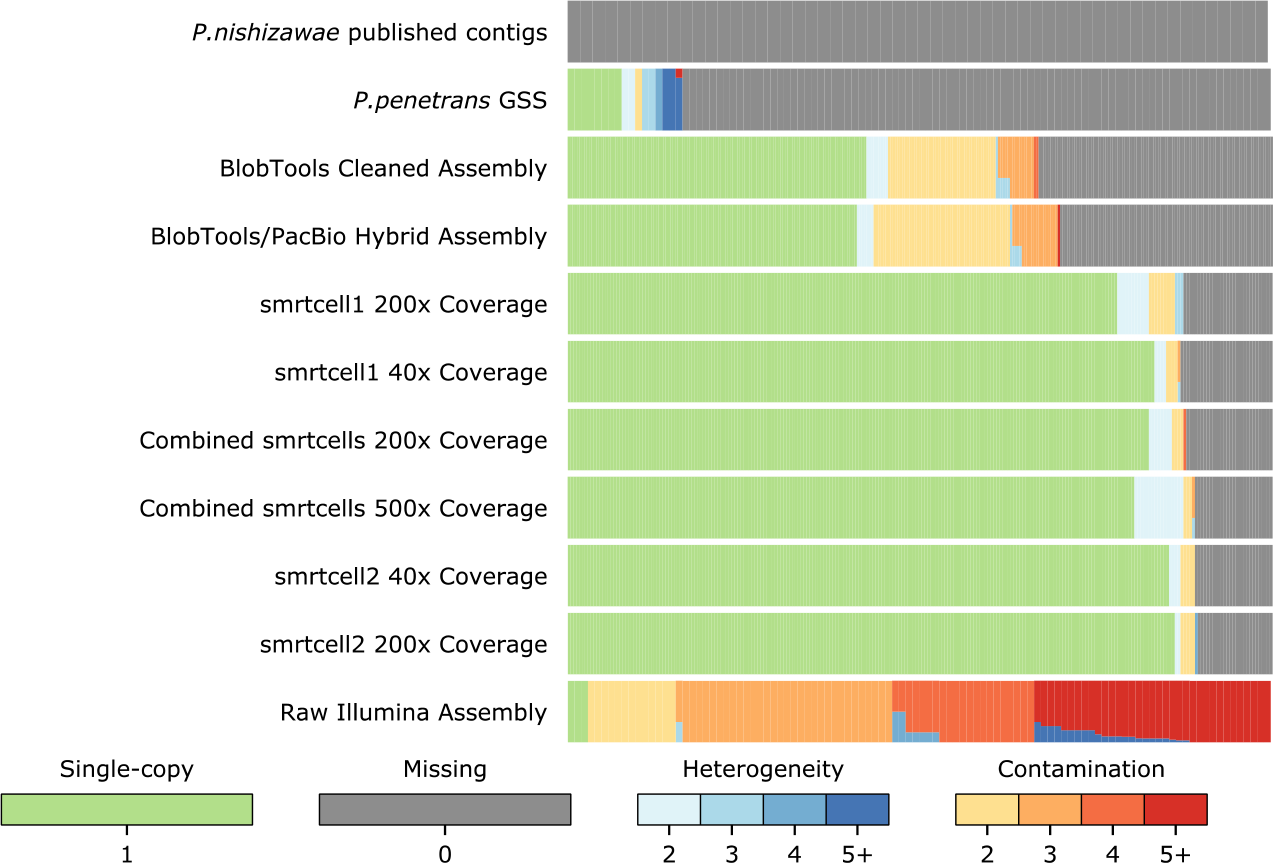
A histogram of genome completeness, heterogeneity, and contamination as assessed by the presence and length of lineage specific marker genes for various *Pasteuria* assembly versions. This figure was created using bin_qa_plot in CheckM

**Figure 2:**
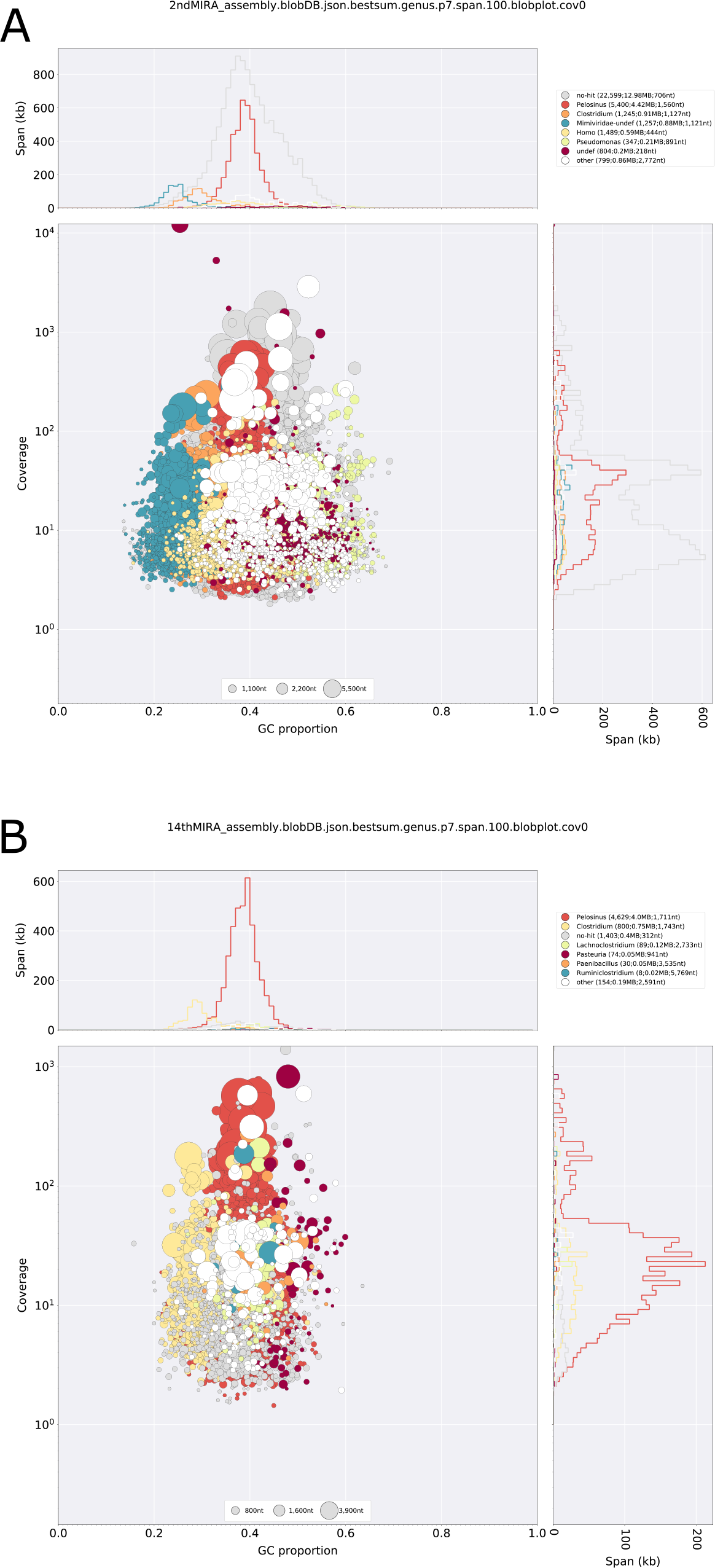
BlobTools GC coverage plot of MIRA assembled legacy Illumina reads after two (a) and 14 (b) iterations of contaminant read removal with taxonomic assignment from BLAST hits at the genus level.

Contamination and heterogeneity were consistently lower in PacBio only assemblies; while completeness was typically higher, except for raw Illumina assemblies whose completeness score was inflated by contaminant markers. Hybrid assembly of the raw or BlobTools cleaned Illumina reads with initial SMRT cell long reads offered a slight improvement on either Illumina assembly but a significant decrease in the overall quality of the same PacBio data assembled alone.

Comparison with existing published genomic sequences revealed high identity alignment with our PacBio assembly (Fig. 3a), although the coverage and length of alignments was often limited (Fig. 3b). Of the 2.4Mbp genome survey sequence (GSS) [17] 0.48Mbp aligned with our genome with 98.5% identity. Legacy Illumina data, which had been restricted to firmicute contigs using the BlobTools pipeline, aligned with 99.4% identity to 0.77Mbp of our assembly. In contrast, 1.97Mbp of the legacy Illumina assembly aligned with 95% identity to the *Pelosinus fermentans* genome. ANIm of the published *P. nishizawae* contigs aligned to only 286bp of both *P. penetrans* PacBio assembly and GSS sequences with 88.5% identity.

**Figure 3:**
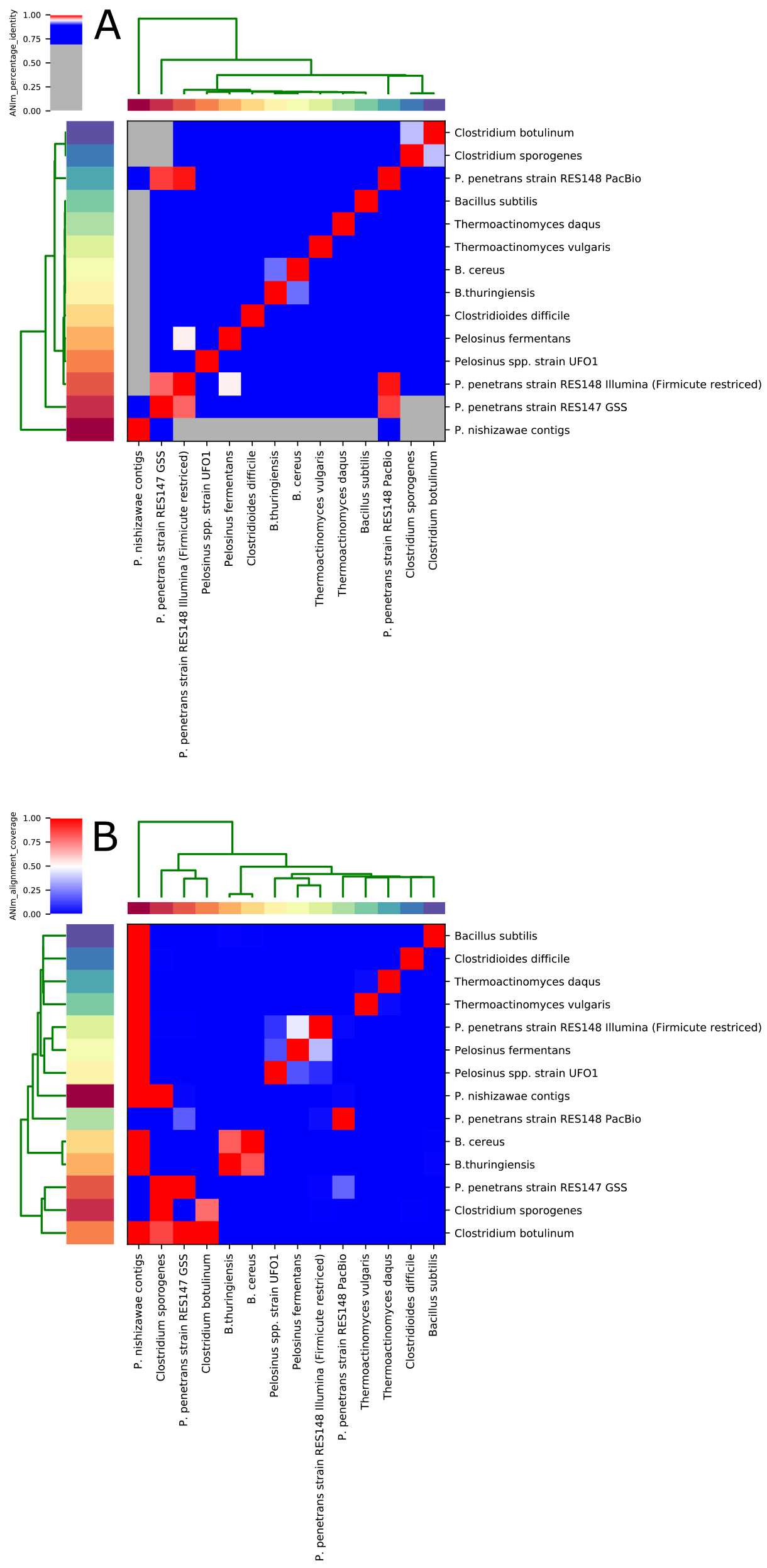
ANIm alignment percentage identity (a) and coverage (b) of PacBio and legacy illumina assemblies with related firmicutes and published *Pasteuria* spp. sequences.

### 6.2 Comparative Genomic Analysis

Multiple marker gene phylogenetic analysis places *P. penetrans* within the Bacilli. Furthermore, within the Bacilli *P. penetrans* is most closely related to *Thermoactinomycetae* (Fig. 4).

**Figure 4:**
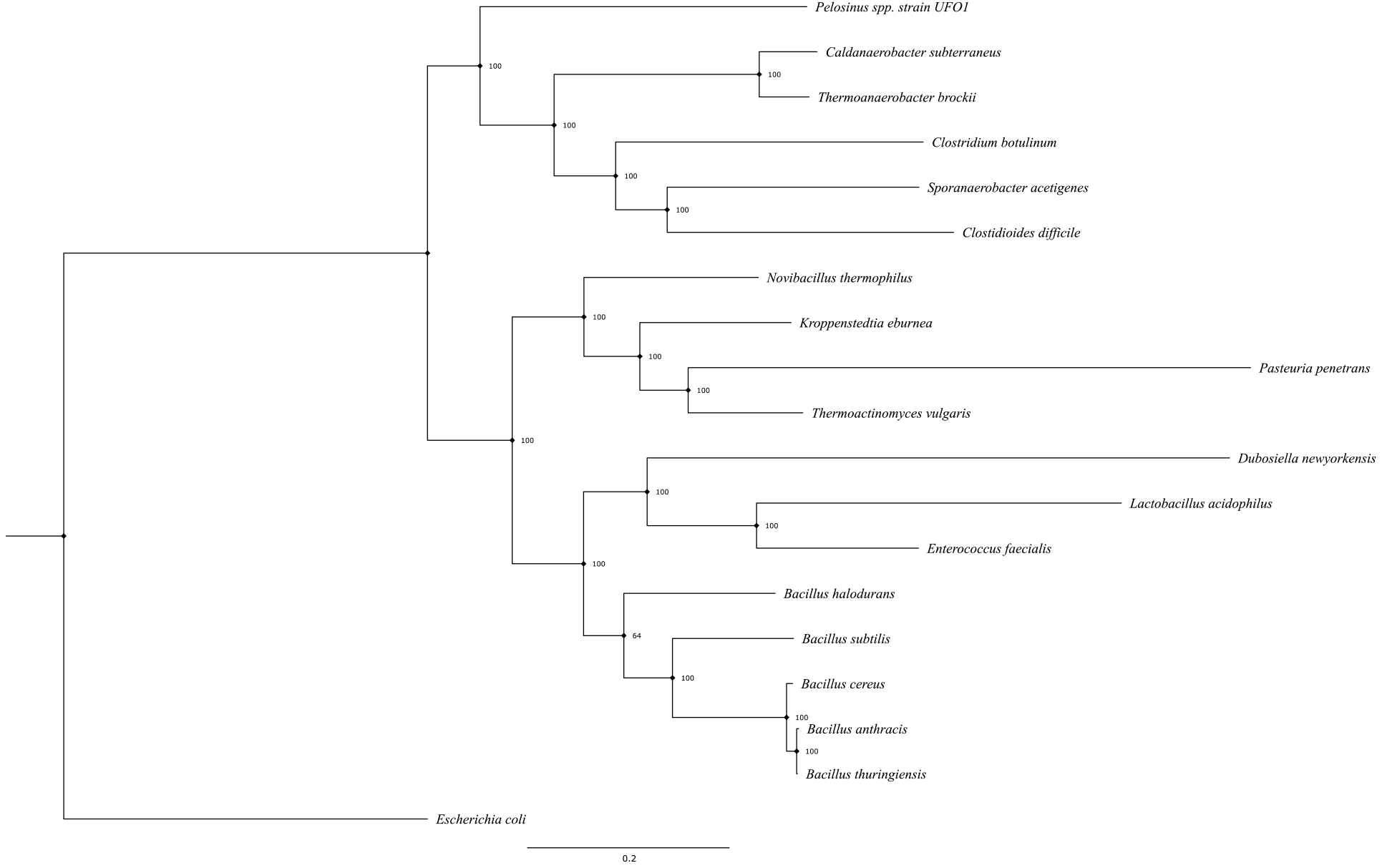
Multi-gene maximum likelihood phylogeny of *Pasteuria penetrans* within Firmicutes. Branches are labelled by bootstrap support. Produced in bcgTree and graphically represented within figtree. 109 essential genes are aligned from provided proteomes to generate this tree.

*Pasteuria penetrans* contained the most unique clusters both in absolute and relative terms compared to firmicute genomes included in our analysis (Fig. 5a). Sporulation associated clusters showed much higher conservation (Fig. 5b). *Pasteuria penetrans* contained predicted proteins which clustered with Spo0F, Spo0B, and Spo0A from *Bacillus* species. Spo0A and Spo0F were also annotated by BlastKOALA; Spo0B was not. No SinI or SinR domain containing proteins were predicted from *P. penetrans*.

**Figure 5:**
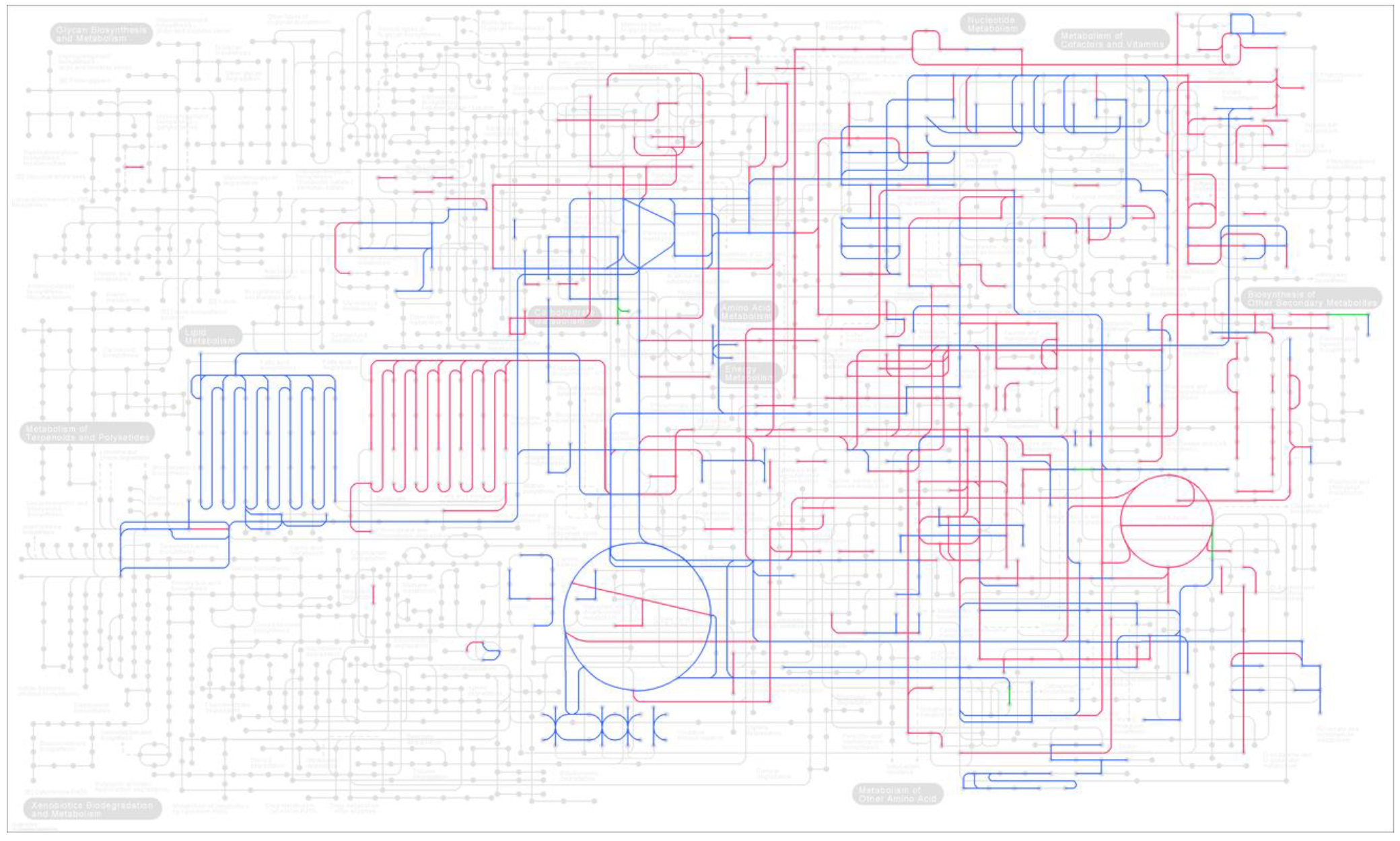
KEGG reconstruct pathway metabolic overview map of *P. penetrans* and *Thermoactinomyces vulgaris* KOALABlast output. Pathways predicted in both organisms are coloured in blue; pathways predicted in *T. vulgaris* only in red; and in *P. penetrans* only in green.

Of 3511 unique *P. penetrans* protein clusters 136 were annotated with transposase domains, 15 with collagen triple helix domains, and 3223 were not annotated. An additional two transposase protein clusters were shared by *P. penetrans*, *B. thuringiensis*, and *T. vulgaris*, giving a total transposase cluster count of 138 in *P. penetrans*. The total number of transposase annotated clusters was 163 across all predicted proteomes. One *P. penetrans* protein functionally annotated with a collagen triple helix repeat clustered with six proteins of *B. thuringiensis* and three proteins of *C. difficile*.

### 6.3 Metabolic modelling

*Pasteuria penetrans* showed a reduced metabolism relative to *Thermoactinomyces vulgaris* (Fig. 6), returning 755 KEGG orthologues compared to 1871, representing a relative reduction of 59.6% in components of well characterised pathways. The reduction of *P. penetrans* genome size is approximately 30% relative to *T. vulgaris*.

**Figure 6:**
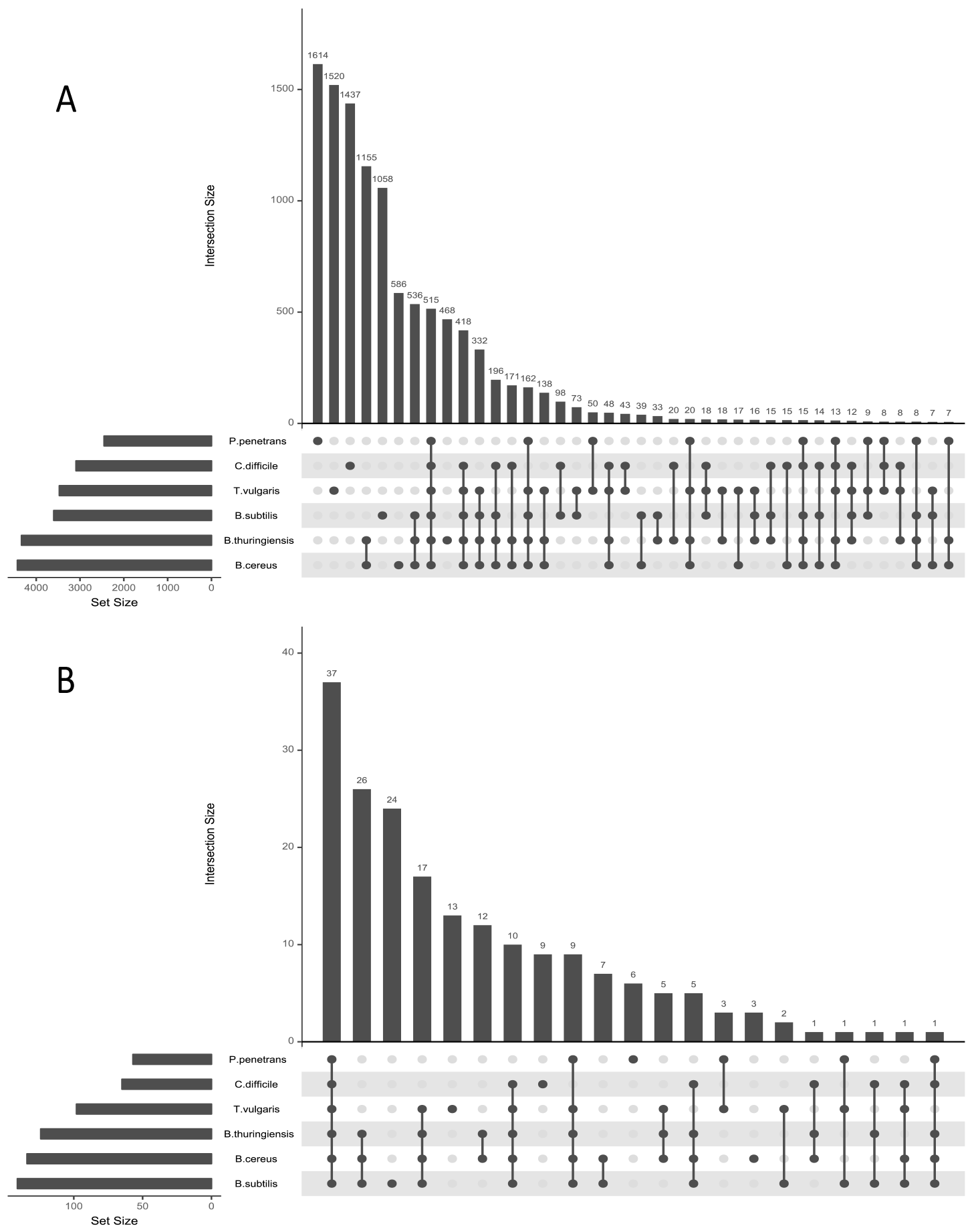
(a) an intersect plot produced in UpSetR showing clusters shared between proteomes indicated by connecting dots below the x axis and ordered by total number of clusters. (b) An intersect plot of clusters which returned sporulation or spore related terms from InterProScan functional annotation.

When compared to the plant parasitic nematode symbiont *Xiphinematobacter* spp. and the *Wolbachia* symbiont of the filarial parasite *Brugia malayi* (*wBm*), *P. penetrans* showed a comparative reduction in pathways, each of these returning 572 and 545 KEGG orthologues respectively.

*Pasteuria penetrans* appears to possess a complete fatty acid biosynthesis pathway, although lacks the fatty acid degradation pathway in its entirety. Both *wBm* and *Xiphinematobacter* spp. also lack this pathway. Enzymes involved in glycolysis are absent up to and including the conversion of alpha-D-glucose 6-phosphate to beta-D-fructose 6-phosphate. Similarly, the pentose phosphate pathway includes no glucose processing enzymes appearing to begin at β-D-fructose 6 phosphate and/or D-ribulose 5 phosphate. *Pasteuria penetrans* also possesses a partial chitin degradation pathway capable of degrading chitin to chitobiose and N-acetyl D glucosamine.

Synthesis pathways for a significant majority of amino acids are absent except for Aspartate and Glutamate. Conversion of glycine to serine and vice versa is predicted due to the presence of *glyA*. The lysine biosynthesis pathway proceeds only as far as miso-diamelate which feeds directly into a complete peptidoglycan synthesis pathway. Purine and pyrimidine biosynthesis pathways are present but appear to be peripherally reduced. Several predicted proteases are also present.

ABC transporters carrying zinc, iron (II), manganese, phosphate, and branched chain amino acids are present. An additional nucleotide binding ABC transporter implicated in cell division and/or salt transport is also present. One component of an Iron complex transporter (FhuD) is predicted. From this model, isoprenoid biosynthesis appears to proceed following the non-mevalonate pathway. No pathways for the biosynthesis of siderophores were predicted from this assembly. None of the components of a flagellar assembly were observed.

Sec-SRP and Twin arginine targeting (TAT) secretion pathways are predicted from KEGG orthologies. We did not find evidence of orthologues to characterised toxins or virulence factors in the *P. penetrans* genome.

A complete pathway for prokaryotic homologous recombination is predicted in our assembly. Base excision and mismatch repair machinery also appears to be intact. Competence related proteins ComEA and ComEC are predicted from KEGG orthologues. KinFin analysis also returned a putative *P. penetrans* orthologue for ComEA as well as predicted proteins which clustered with competence related proteins CinA and MecA from related firmicutes.

### 6.4 Characterisation of collagenous fibres

Collagen domains were identified in 32 unique predicted genes across assemblies of the second PacBio SMRT cell at 40X and 200X coverage. Of these 32: 17 were predicted in both versions of the assembly; 5 were unique to the 40X coverage assembly; and 10 were unique to the 200X coverage assembly.

RaptorX server structural predictions returned significant alignments to BclA/C1q-like structures in 17 of 32 predicted collagens (supplementary text file 2). Fifteen of these consisted of N-terminal collagenous filament domains of varying length each with a C1q/BclA-like C terminal globular head (Fig. 7). The remaining two possessed this predicted structure but contained three and six domains in total. Net charge, Sec/TM domain prediction and binding site predictions for each BclA-like collagen are listed in Table 1.

**Figure 7:**
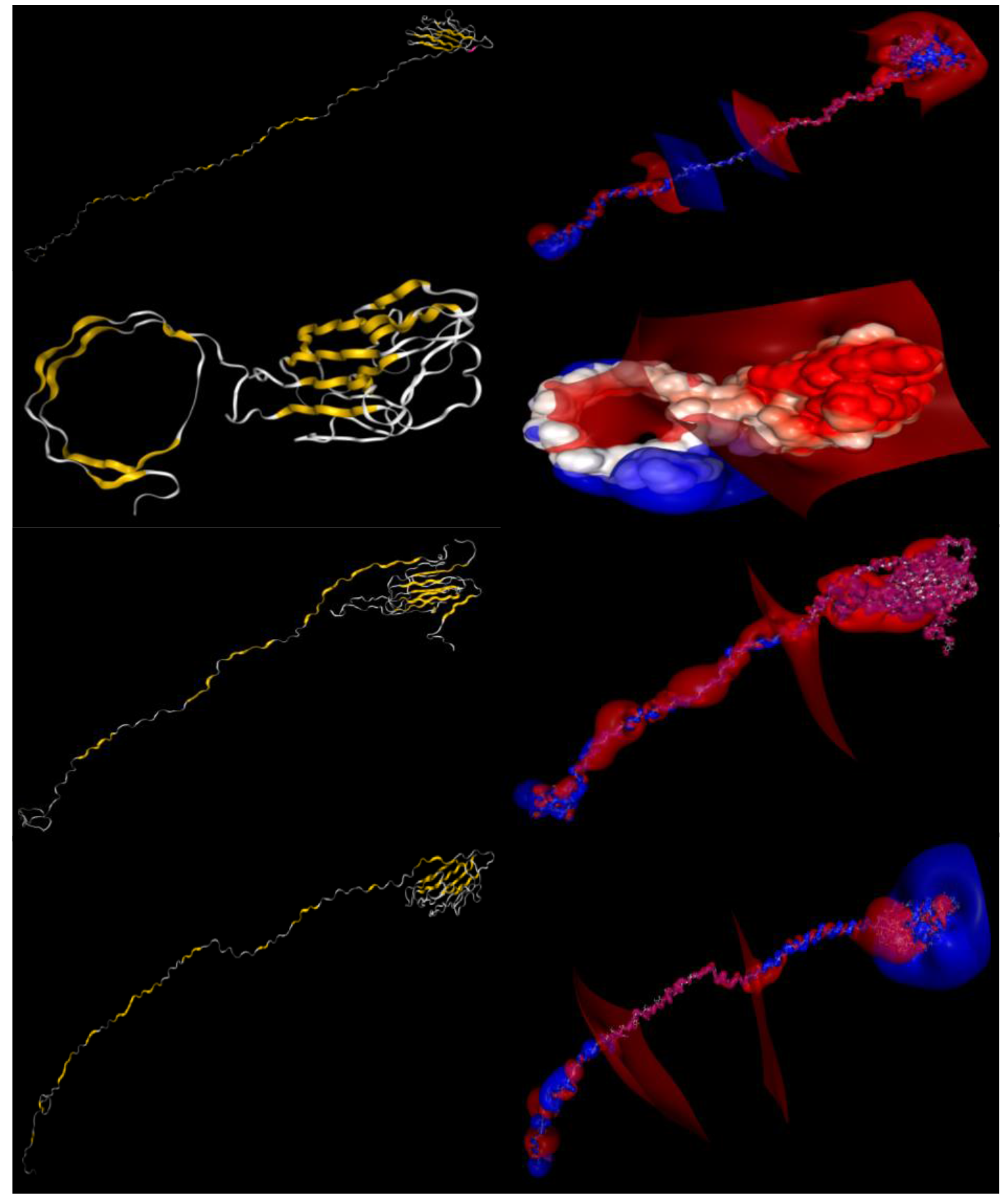
Predicted structure of four BclA-like attachment candidate proteins recovered from the *Pasteuria penetrans* genome. Molecular structure left and corresponding electrostatic surface potential right. Protein structure was modelled in the RaptorX server and electrostatic potential was calculated using the pdbtopqr server and apbs (v1.5). Images were produced using the NGL viewer.

**Table 1:** BclA/C1q-like collagens identified in the P. penetrans genome with net charge from pdb2pqr, Sec/TM domain prediction from PREDTAT, and predicted binding sites in the globular C-terminal domains from the RaptorX server.

| **Collagen** | **Net charge**<br>**(pH=7)** | **Sec/TM**<br>**domain** | **BclA/C1q binding sites** |
| --- | --- | --- | --- |
| **PclP01** | -3 | Sec | NAG(x3), CA, ACT, GOL |
| **PclP02** | -6 | Sec | NAG, EDO(x4), CL(x3), GOL, ACT |
| **PclP03** | 0 | Sec | None |
| **PclP04** | 0 | None | EDO(x3), SEP, U, ACT(x3), IOD |
| **PclP05** | 0 | Sec | None |
| **PclP06** | -2 | Sec | NAG(x2), GOL, DIO(x2), MG, ACT, EDO,<br>144, CL |
| **PclP07** | 9 | None | NAG(x2), CA |
| **PclP08** | -2 | Sec | None |
| **PclP09** | -2 | TM | NAG, DIO(x2), CA |
| **PclP10** | -4 | Sec | NAG(x3), CA, SO4 |
| **PclP11** | -1 | Sec | NAG(x2) GN1, CA |
| **PclP12** | -5 | Sec | NAG(x3), CA, EDO(x2), CA, SO4, DIO, GOL |
| **PclP13** | 0 | None | CA, NAG(x3), EDO(x2), CPS, DIO |
| **PclP14** | 15 | Sec | CA, GN1, NAG(x2), DIO(x3), GOL, CL |
| **PclP15** | -2 | None | CA, NAG(x2), GOL, EDO(x3), CPS, SO4, ACT |
| **PclP16** | 0 | TM |  |
| **PclP17** | -2 | Sec | CA, CL(x2), GN1, NAG, GOL(2), ACT, EDO,<br>GOL SO4 |

Of the 17 previously reported *P. penetrans* RES148 collagen-like proteins [48], one exact match and four additional BLAST alignments at or above 90% identity were found with the 32 predicted collagens identified in our analysis. Of these, two returned a predicted BclA-like structure. No significant alignment was observable between the predicted BclA-like collagens in this assembly and those described in *P. ramosa* [49, 50].

## 7. DISCUSSION

### 7.1 Comparison with legacy assembly data

PacBio only assemblies were of higher quality than hybrid or merged PacBio-Legacy Illumina assemblies. The high identity of marker gene BLAST hits, coupled with the presence of multiple distinct marker gene copies, and clear separate peaks on GC coverage plots indicate true contamination in the legacy Illumina dataset as opposed to *P. penetrans* alignment to related firmicutes. Marker gene alignments, GC coverage plots, and ANIm alignments point to significant *Pelosinus* spp. contamination in the firmicute restricted legacy Illumina data assembly. Initial GC coverage plots also point to contamination from Mimiviridae, human, *Clostridium* spp., and *Pseudomonas* species in the unrestricted assembly analysed by Srivastava *et al.* [48].

The genome size of *P. penetrans* has been estimated to be between 2.5 and 4Mbp with a GC content approximately similar to that of *Bacillus subtillis* and *B. halodurans* at 44% [51]. Our assemblies are consistently placed at the lower end of this size range, with a GC content of around 46%. An unpublished *P. penetrans* genome approaching 2.5Mbp was described by Waterman *et al.*, [18] however, as this data has not been made available it is not possible to evaluate this assembly directly. Our assembly is small with reference to free living bacilli but large in comparison to other bacteria obligately associated with nematodes such as *wBm* (~1.1Mbp) [52] and *Xiphinematobacter* spp. (~0.9Mbp) [53]. The completeness score of our assembly was high at 86% based on lineage specific marker genes. Notably, the same lineage specific markers were not predicted in PacBio assemblies at varying levels of coverage. This may indicate the interference of sequencing or amplification errors in gene prediction.

### 7.2 Phylogeny

Maximum likelihood phylogenetic analysis of core genes (Fig. 3) confirms the position of our sequence within the endospore forming Bacilli with strong bootstrap support [51, 54]. However, it was not possible to determine that *Pasteuria* spp. are ancestral to *Bacillus* spp. as previously described [51]. Early observations of *Pasteuria* spp. pointed to a potential grouping with *Thermoactinomyces* based on morphological comparisons, noting that both *P. penetrans* and *T. vulgaris* form filamentous tubes on germination as opposed to vegetative rods [2]. These morphological comparisons were also observed by Ebert *et al.* [55], however, these researchers, and many others thereafter, highlight that genetically inferred phylogenies point to a more distant relationship than these morphological similarities might suggest [9, 18, 51, 54, 55]. Despite the apparent distance of this relationship *Thermoactinomyces* spp. remain the closest observable relations within Bacilli within our analysis. Genomic sequencing of *Thermoactinomyces* sp. strains AS95 and Gus2-1 display similarly small genomes of 2.56Mbp and 2.62Mbp respectively with GC content around 48% [56, 57]. Both *Thermoactinomyces vulgaris* and *Thermoactinomyces daqus* H-18 were however found to be larger at 3.70Mbp and 3.44Mbp respectively [58, 59].

### 7.3 Plasticity

The largest component of *P. penetrans* specific predicted proteome clusters which returned InterPro annotations contained transposase domains. This is surprising as transposases typically constitute a lower proportion of genes in smaller genomes generally, and are completely absent in most obligate, host-restricted bacteria [60, 61]. However, enhanced genome plasticity may be better tolerated by bacteria which have recently adapted to a symbiotic or pathogenic lifestyle which are able to compensate for non-specific transposase insertions due to functional redundancy enabled by the host [62, 63]. Indeed, genome plasticity may be selected for in such organisms allowing for faster adaptation to the host [61]. This does not necessarily indicate the recent conversion to obligate parasitism in the case of *P. penetrans* as some ancient symbiotic bacteria, such as *Wolbachia pipientis*, may also exhibit high numbers of transposable elements (TEs) [64]. This may be explained by the “intracellular arena hypothesis” where foreign TEs exchanged during a host switching event are retained because they are advantageous or are better tolerated by obligate bacteria [65]. Kleiner *et al.*, described high transposase gene number and expression in symbiotic bacteria of the oligochaete worm *Olavius algarvensis* [61]. They hypothesised that loss of tight regulation of transposase expression may play a role in the expansion of TEs in host-restricted bacteria and thus in their adaptation to the host. Notably, the genomes of the tropical apomictic RKNs, from which *P. penetrans* was isolated, also exhibit extensive plasticity [66–70] thought to promote their extraordinarily broad host range, compared to their sexually reproducing counterparts, which encompasses most flowering plants [71].

In addition to transposase expansion *P. penetrans* appears to possess complete bacterial homologous recombination pathways and partial components of known competence pathways. Waterman [18] described the presence of competence related ComC, ComE, and ComK predicted proteins from their *P. penetrans* genomic assembly. The presence of ComEA, ComEC, CinA, and MecA predicted orthologues supports their assessment of the potential of *P. penetrans* for competence although we did not find ComC, or ComK in our assembly. RecA, which is involved in DNA repair, recombination, and competence [72] was also present in both assemblies.

### 7.4 Metabolic pathways

The reduction of metabolic pathways does not scale with the reduction in total genome size with the reduction in genome size when compared to *T. vulgaris* being approximately half the reduction of known metabolic pathways. *Pasteuria penetrans* shows a reduction of metabolic pathways which is comparable to *Xiphinematobacter* spp. and *wBm* despite a total genome size more than double both organisms [52, 53]. The large number of unannotated gene predictions which do not cluster with related proteomes may suggest the use of alternate pathways.

Also notably absent are synthesis pathways for the majority of amino acids again mirroring the metabolism of *wBm* [52] and to a lesser extent *Xiphinematobacter* [53]. This, combined with the presence of a branched chain amino acid transporter and multiple proteases, suggests that *P. penetrans* is near completely reliant on the host for amino acids.

Less clear is where *P. penetrans* may be acquiring carbon. The absence of glucose, sucrose, mannose, and starch catabolising pathways in the assembled genomic sequence is notable. This parallels the metabolic profile of *wBm* which rely on pyruvate dehydrogenase and TCA cycle intermediates produced by the degradation of proteins [52]. Possible carbon sources include fructose, and/or partial gluconeogenesis from TCA cycle intermediates such as citrate, malate, fumarate, and succinate. Initial D-fructose phosphorylation to fructose-1-P is not predicted but complete pathways from conversion of fructose-1P to both glyceraldehyde-3P and fructose-6P appear to be present.

Duponnois *et al.* [15] identified an apparent positive influence of *Enterobacter* spp. on the development of *P. penetrans* in the field. Production of organic acids in the rhizosphere by *Enterobacter* spp. is well documented [73] and the production of such acids in *Enterobacter* spp. culture filtrates is highlighted in *in vitro Pasteuria* spp. culture patents filed by Gerber *et al.* [74]. Earlier unsuccessful attempts to culture *P. penetrans in vitro* also noted that culture filtrates from *Thermoactinomyces* spp. and fungi were capable of improving the maintenance of *P. penetrans* replicative stages [14]. *Bacillus subtillis* is capable of growing with citrate as a sole carbon source [75], whilst iron citrate uptake is required for the virulence of *B. cereus* [76].

*Pasteuria penetrans* appears capable of breaking chitin down into NAG and chitobiose. Possible functions of these enzymes are in the degradation of chitin as the structural component of nematode eggs or in the breakdown of mucins in the nematode cuticular matrix. Simple digestion of the nematode egg shell may explain the mechanism by which *P. penetrans* reduces host fecundity. The apparent involvement of NAG in attachment may provide another important requirement for chitin catabolism.

### 7.5 Collagen-like proteins

Of 32 unique collagens identified by pooled gene prediction algorithms 17 returned structural alignments matching C1q or BclA like proteins. Collagenous fibres on the surface of endospores within the bacilli are often components of the infection process. Among these the most well characterised is BclA, a C1q-like collagenous glycoprotein which forms the hair-like nap of fibres present on *B. anthracis* [77]. C1q is a component of human complement pathway which binds IgG, and apoptotic keratinocytes [78, 79]. BclA is implicated in a number of processes in *B. anthracis* infection including specific targeting to macrophages [80], and immunosuppressive activity via binding to complement factor H [81]. Notably, BclA does not appear to be directly involved in attachment as ΔbclA spores show no reduced binding to host cells but do exhibit a reduction in specific targeting to professional phagocytic cells [80]. Conversely, three fibres paralogous to BclA are also described in *C. difficile* which appear to be directly involved in the early stages of infection [82, 83]. Further, it has been demonstrated that *C. difficile bclA^−^* mutants display reduced adherence to human plasma fibronectin [84]. Fibronectin has been proposed as a binding target of *Pasteuria* spp. spores [85]. However, fibronectin does not appear to be an abundant component of the *Meloidogyne incognita* J2 cuticle [86].

Along with this set of BclA-like collagens, CotE is also predicted from our assembly where it is possible that it might similarly be involved in a multi-component attachment process. The CotE protein is also thought to be involved in the colonisation of the gut by *C. difficile* through C-terminal binding and degradation of mucins [87]. Glycosylated mucins on the nematode cuticle are implicated as the target in the ‘Velcro’ model of attachment [12].

Variation in the lengths of the predicted collagenous fibres reflects the observable variation in endospore surface fibres observable with electron microscopy [12]. The presence of 17 such fibres may be indicative of functional redundancy. It has been observed that NAGase treatment can invert endospore attachment in some instances [9] and in cross-generic attachment of *Pasteuria* HcP to *Globodera pallida* endospores are predominantly inverted in their attachment to the cuticle [88]. The absence of the majority of the collagen sequences reported by Srivastava *et al.* [48] likely reflects the contamination we report within the legacy Illumina data assemblies. No alignment to putative collagens identified in *Pasteuria ramosa* [49, 50, 89, 90], infective of the water flea *Daphnia magna*, was observed. If specificity is determined by the presence of unique collagens, or unique sets of collagens, then this is unsurprising due to the vastly different host range of these species. Further sequencing across species or strains of this genus with differing attachment profiles may reveal divergent BclA-like collagen profiles. The nature of attachment is of practical interest to the application of *Pasteuria* spp. as biocontrol agents; however, in the case of *P. ramosa*, elucidation of the molecular mechanics of this interaction may impact our fundamental understanding of host-parasite evolution as this infection system is a prominent model for the study of the red-queen hypothesis [49, 50, 55, 89–92]. The variable electrostatic potential of these predicted proteins is notable as it has been suggested that electrostatic interactions may play an initial role in the attachment process; the electrostatic potential of *P. penetrans* endospores having previously been characterised as negative [93]. The predicted collagens in this assembly match very well with our expectations of the molecular components of attachment based on experimental evidence to date. However, further work is required to evaluate their role in this process, and to evaluate the Velcro-like attachment model.

## Supporting information

## 9. ABBREVIATIONS

GSS: Genome Survey Sequence
MDA: Multiple Displacement Amplification
WGA: Whole Genome Amplification
NAG: N-acetyl-D-Glucosamine
RKN: Root Knot Nematode
WBm: Wolbachia of Brugia malayi
SDS: Sodium Dodecyl Sulphate
TCA: Tricarboxylic Acid
TE: Transposable element
ANI: Average Nucleotide Identity

