## Supplementary material for "*De novo* assembly of the *Pasteuria penetrans* genome reveals high plasticity, host dependency, and BclA-like collagens"

Supplementary methods file 1.

**Whole genome amplification.**

The illustra GenomPhi V2 DNA amplification kit (GE Healthcare) was used with *P. penetrans* DNA template as per manufacturer’s instructions. 10 ng of template DNA was used per reaction however isothermal amplification at 30 ^o^C was allowed to proceed for 3 h before reactions were stopped. A total of 8 individual reactions were prepared and pooled for each template type, after completion of isothermal amplification. A 1 µl aliquot of each WGA reaction pool was subjected to electrophoresis with 1 % agarose gels and subsequently visualised by staining with ethidium bromide (0.5 *µ*g ml^-1^) to determine whether genomic amplification had been successful.

**phi29 debranching of WGA DNA.**

Before debranching a mixture of 8 WGA reactions were column purified with the Min Elute kit (Qiagen) into 40 µl of dH_2_0. Samples were debranched in a total of 50 µl with 10 U phi29 (Fermentas), 10 X buffer phi29 reaction buffer (Fermentas), 1 mM dNTPs (Fermentas) and incubated at 30 ^o^C for 3 h. Reactions were terminated by heating at 65 ^o^C for 10 min.

**S1 nuclease treatment.**

Prior to S1 nuclease treatment the phi29 treated sample was column purified and resuspended in 165 µl dH_2_O. This was added to 200 U S1 nuclease (Fermentas) with 5 X reaction buffer. The mixture was heated at 37 ^o^C for 30 min, after which time the enzyme was deactivated by addition of 100 µl of 100 mM EDTA and heated at 70 ^o^C for 20 min. Finally, the sample was column purified and resuspended in 50 µl dH_2_O.

**Preparation of sample for Illumina^®^ sequencing and assembly.**

The *Pasteuria penetrans* sample was sequenced at The Genome Analysis Centre, Norwich, UK on Illumina^®^ GAIIx with a paired end run with read length of 120 bp. These reads were then assembled using AbySS-1.2.7 [1], with a kmer length of 61.
